# Benchmarking cellular segmentation methods against Cellpose

**DOI:** 10.1101/2024.04.06.587952

**Authors:** Carsen Stringer, Marius Pachitariu

## Abstract

In a recent publication, Ma et al [1] claim that a transformer-based cellular segmentation method called Mediar [2] — which won a Neurips challenge — outperforms Cellpose [3] (0.897 vs 0.543 median F1 score). Here we show that this result was obtained by disadvantaging Cellpose in multiple ways. When we removed these impairments, Cellpose outperformed Mediar (0.861 vs 0.826 median F1 score on the updated test set). To further investigate the performance of transformers for cellular segmentation, we replaced the Cellpose backbone with a transformer. The transformer-Cellpose model also did not outperform the standard Cellpose (0.848 median F1 test score). Our results suggest that transformers do not advance the state-of-the-art in cellular segmentation.

---

Automated cellular segmentation has advanced substantially in recent years. Our own efforts in this direction have resulted in Cellpose, an easy-to-use segmentation algorithm that biologists have found useful [3]. Cellpose has become a standard against which new cellular segmentation algorithms are benchmarked. For example, “foundation” models that use transformer architectures have recently claimed substantial progress versus Cellpose [1, 2, 4, 5]. However, the comparisons in these papers may not always be conducted under appropriate conditions. Thus the question often arises: “does algorithm X outperform Cellpose?”. Here we investigate the benchmarks in the challenge proposed by Ma et al [1] and find that the Cellpose model used for comparison was disadvantaged in multiple ways. When trained appropriately, Cellpose with default parameters achieves state-of-the-art performance on this dataset.

## Challenge details

Ma et al [1] reports on the results of a cellular segmentation competition held at Neurips 2022 in which a method called Mediar won [2]. This competition had the same general goal as the original Cellpose study: to produce methods that work out-of-the-box on a wide range of cellular images. To this purpose, the organizers generated a new annotated training set of 1,000 images from various modalities, with an additional 101 images for validation and 400 images for testing. These images roughly covered the same modalities as those in the original Cellpose dataset, with some modalities covered better (i.e. phase-contrast, DIC) and other modalities covered less well (fluorescent). Separately from the challenge, the authors shared comparisons with established methods including Cellpose. These comparisons appeared to show that the methods developed for the challenge make a gigantic leap in performance over Cellpose (median F1 score increased from 0.543 to 0.897 on average).

The poor performance of Cellpose was unexpected, because the images from the benchmark *look* like images that Cellpose should be able to segment (Figure 1a,b). In addition, the top performing algorithm in the challenge, Mediar [2], uses the Cellpose framework [3], with the same flow field representation combined with pixel class prediction for training; the same flow field dynamics used to reconstruct cells from the predictions; and the same quality control step using the error between predicted and reconstructed flow fields. In fact, Mediar directly copied our codebase for implementing the Cellpose framework without modification.

**Figure 1.**
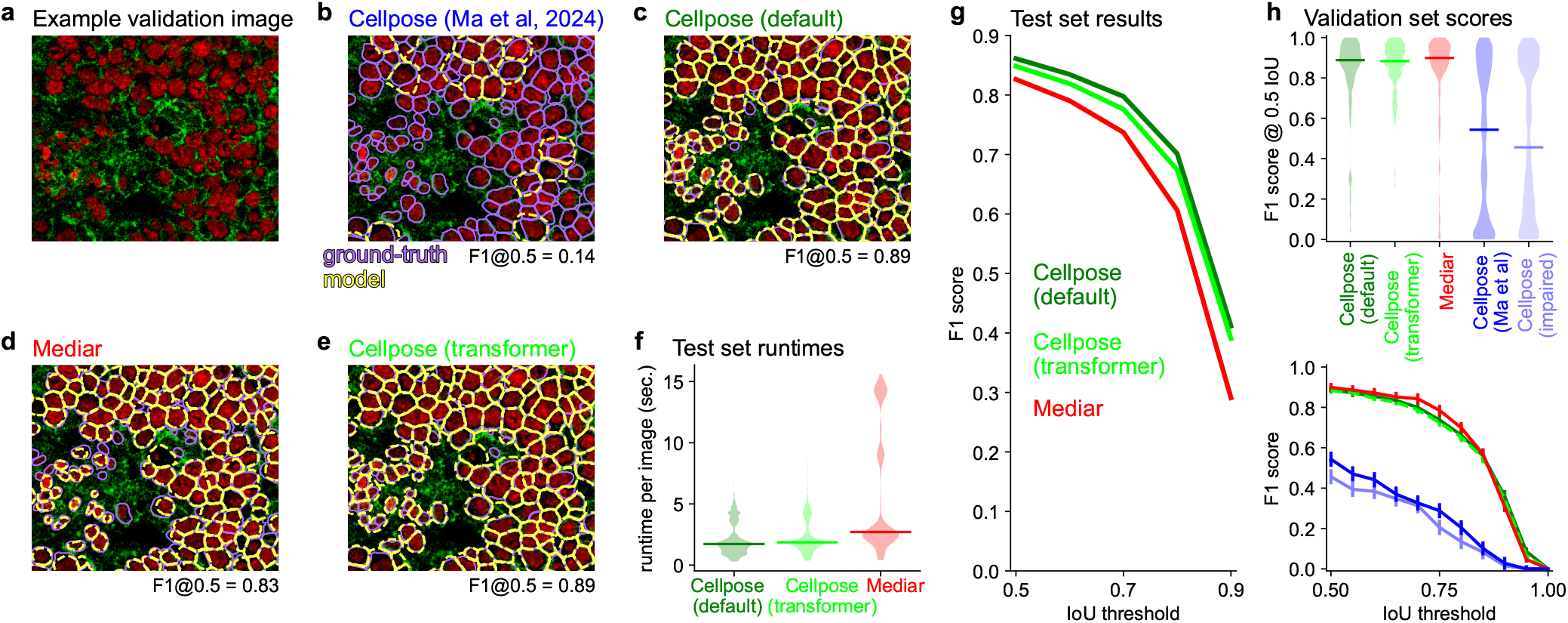
Cellpose trained appropriately performs well. **a**, Example image from the validation set. **b**, Same image with ground-truth segmentation overlaid in purple, and segmentation from Cellpose model from Ma et al [1] in yellow. **c**, Same as **b**, for the default Cellpose model trained without disadvantages. **d**, Same as **b**, for the Mediar model [2]. **e**, Same as **b**, for the Cellpose model with a transformer backbone. **f**, Per image runtimes for each algorithm on the 400 test set images, median shown as a solid line, on an A100 GPU. **g**, Median performance across the 400 test set images, for each algorithm at different intersection over union (IoU) thresholds. **h**, Top: F1 score at 0.5 IoU threshold for the 100 validation images across all algorithms, median shown as a solid line. Bottom: Median F1 score for the 100 validation images across IoU thresholds, error bars represent s.e.m.

Ma et al [1] does not dwell on this similarity between Mediar and Cellpose. Instead, they emphasize the novelty of using a transformer architecture for prediction, and claim that the new architecture provides the observed massive boost in performance over Cellpose. The comparisons, as we show below, are to a version of Cellpose that has been artificially disadvantaged across multiple axes.

### Cellpose configuration in Ma et al [1]

To compare the performance of machine learning algorithms, it is essential to use a consistent benchmark methodology for training and testing. Otherwise, it cannot be known whether observed differences in performance are due to differences between the algorithms or due to differences in methodology. Several such differences stand out in the training and testing of Cellpose from Ma et al [1], which have detrimental effects on the Cellpose results.

*Disadvantage One*. Cellpose was trained and tested on grayscale images while the other methods were trained on the full multi-channel data. Clearly, multi-channel images contain substantially more information. The channels could have been fed into Cellpose in their original order, and Cellpose would have dealt with this in the same way as the top algorithm does (which uses the Cellpose framework for mask prediction). Note that despite being run on grayscale images, the output of Cellpose is illustrated on color images in [1].

*Disadvantage Two*. Cellpose was trained with images rescaled so that cells have an average diameter of 30 pixels (which is the default), but tested with unresized images in which cells have diameters up to 400 pixels. The full Cellpose pipeline includes training a size model for cell diameter prediction at test time, which was not done in Ma et al [1]. This likely explains why training Cellpose on the challenge dataset actually *hurt* its performance compared to the “pretrained” Cellpose which never saw the challenge images [1]: the pretrained model contains a size model, whereas the retrained model does not.

*Disadvantage Three*. The training set for Cellpose included only the challenge images, while the top performing methods used additional datasets. For example, the Cellpose dataset was included in the training of the top three methods but not in the training of Cellpose.

In addition to these disadvantages, we note that Cellpose was run without test-time augmentations (TTA). The Cellpose framework makes it easy to augment images at test time and we described this in the original study [3]. Mediar also adopted the same test-time augmentation framework. Also, Cellpose was trained with a batch size of 32 instead of the default 8, and this may have had an effect on performance due to the regularization provided by batch normalization at small batch sizes [6].

### Test set results with retrained Cellpose

To make a fair comparison to the top algorithms of the Neurips challenge, we retrained Cellpose after removing the disadvantages mentioned above. We emphasize that this is the default configuration of Cellpose with no parameter changes made to improve the scores. We only made minimal changes to the Cellpose training script to account for three-channel inputs. We did not change the default size and rotation augmentations of Cellpose, nor did we change the way the size prediction model is trained. To further make our models comparable to Mediar, we only used the same external datasets they used for augmentation (Cellpose [3], Omnipose [7], Livecell [8] and DataScienceBowl-2018 [9]), despite the availability of other high-quality datasets which we have previously used for the “cyto3” model [10].

Although we used the Cellpose3 codebase [10], the differences from the Cellpose2 codebase [11] used in Ma et al [1] are minimal: 1) we increased the default number of training epochs from 500 to 2,000 to account for bigger datasets, using the AdamW optimizer [12]; 2) we fixed floating point errors in the flow field computation; and 3) we added the option to weigh images in the training set differently depending on their source dataset.

Starting with example segmentations on validation images, we can see that our default Cellpose model is substantially better than the model trained in Ma et al [1] (Figure 1b,c). The segmentations appear similar to those of Mediar, as well as to a version of Cellpose where we replaced the core neural network with a transformer (Figure 1d,e, [10]). Cellpose was overall faster than Mediar (Figure 1f), though the runtime of both models is likely much larger than it would be in practical use with smaller images, batched computations, disabled test-time augmentations, and without running the size model for Cellpose.

We then submitted our results on the 400 test images to the challenge website, which was kept live after the challenge ended although with a modified testing set, in which the organizers replaced 50 of the easiest images with harder images. We submitted the Cellpose results exactly once using default parameters. We also submitted the Mediar results in order to obtain their performance on the updated test set. We achieved a test set median F1 score of 0.861 using Cellpose, which was substantially better than the 0.826 score of Mediar (Figure 1g). The difference was even larger at higher IoU thresholds, which measure how well the algorithms track the precise contours of cells (0.412 vs 0.291 at 0.9 IoU for Cellpose vs Mediar). Thus, the winner of the Neurips challenge was in fact outperformed by Cellpose with default parameters.

### Further analyses on the validation set

Since the ground truth of the test data was not shared, we cannot make further comparisons on these images to understand how the models perform. For the remainder of this paper, we turn to the validation set of 101 images for further analyses (Figure 1h). These results should be interpreted with caution because the competition teams obtained repeated feedback from the validation data, thus likely overfitting to the validation set. In contrast, we did not use feedback from this image set to optimize Cellpose, since we used the default Cellpose parameters. We exclude the very large slide image from these comparisons (it was benchmarked inaccurately in Ma et al [1], see Methods).

The default Cellpose model substantially outperformed the Cellpose model from Ma et al [1] (Figure 1h). Furthermore, we can show that the decrease in performance was mainly due to the disadvantages outlined above: when we retrain Cellpose on grayscale images, with mismatched train-test cell sizes, without additional data, without test-time augmentations and with a batch size of 32, we obtain similarly poor results as those in Ma et al [1] (shown in Figure 1h as “Cellpose (impaired)”).

Despite being outperformed by Cellpose with default parameters, Mediar nonetheless outperformed all the other participants in the challenge. This may not be surprising because Mediar uses the Cellpose framework for mask prediction, while replacing the standard CNN with a transformer and adding a complex multi-step training procedure with multiple types of data augmentation and data resampling steps. To further investigate whether the multi-step training procedure in Mediar is beneficial, we ask whether simply replacing the CNN backbone of Cellpose with a transformer would retain the same performance. To do this, we used the same encoder-decoder transformer architecture as the one used in Mediar. We find that transformer-Cellpose performs almost as well as standard Cellpose (Figure 1g,h), similar to the results in [10] and thus better than Mediar. Since the transformer-Cellpose architecture and training objective are very similar to the ones used by Mediar, this suggests that the custom training protocol for Mediar did not improve test set performance.

### Analyzing the images from Ma et al [1]

In this last section, we use the segmentation model we trained to try to understand how the new dataset in Ma et al [1] relates to existing datasets and what makes this dataset difficult for Cellpose and Mediar respectively. We start by looking at some of the validation images where Cellpose and Mediar both performed well, where Cellpose performed better or where Mediar performed better (Figure 2a). Generally, we find that Cellpose performs better on tissue images, while Mediar performs better on DIC images, particularly those similar to training set images where only a subset of cells are labeled. We do not think these are fundamental limitations of either model. Rather, this difference may result from discrepancies in labeling style between different images in the Ma et al [1] dataset relative to labeling styles used in other datasets. As we have previously discussed elsewhere [11], such discrepancies can negatively interfere with each other.

**Figure 2.**
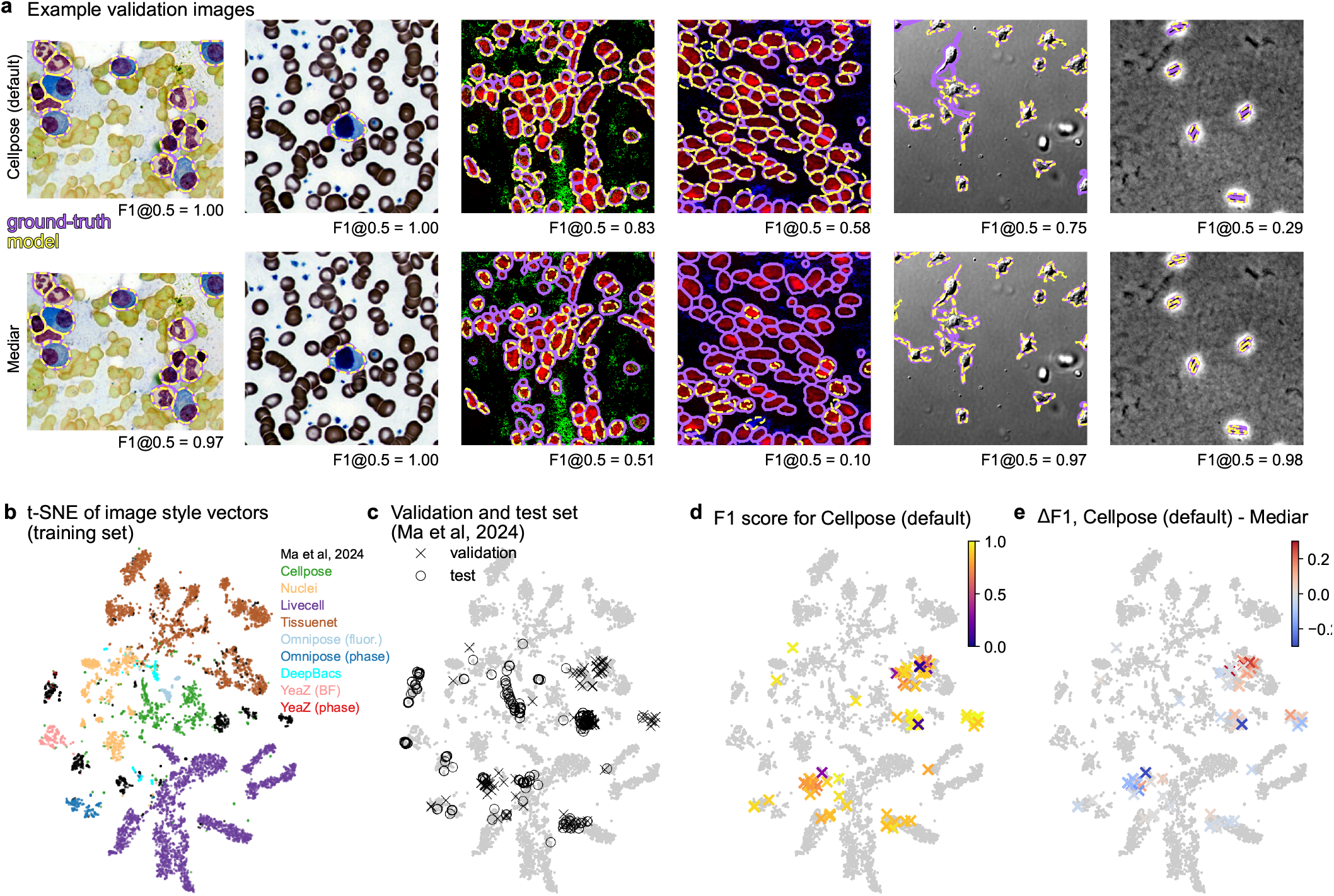
Exploring the dataset of Ma et al (2024). **a**, Example images from the validation set with paired segmentations by Cellpose (top), and Mediar (bottom). Both Cellpose and Mediar performed well on the first two images, while Cellpose performed better on the next two images, and Mediar performed better on the last two images. **b**, Two-dimensional embedding visualization of style vectors from all training set images from Ma et al (2024) together with images from other datasets. **c**, Same as **b** with all images in gray and with validation images from Ma et al (2024) overlaid as black X’s and test images overlaid as black O’s. **d**, Same as **b** with validation images colored by the performance of the Cellpose default model. **e**, Same as **b** with validation images colored by the difference in performance between Cellpose and Mediar.

To get an overview of the images from Ma et al [1] in the context of existing datasets, we performed a two-dimensional embedding of the style vectors from a Cellpose model trained on all available images from different datasets, with 10% of the images coming from the challenge dataset [3, 13]. This visualization clearly groups together images from the same dataset and reveals details in the relative distribution of images from different datasets (Figure 2b). We see that the images from the challenge occupy six relatively tight clusters plus a widely distributed cluster that overlaps well with the Tissuenet dataset. Focusing on the validation and test set of the challenge, we see that only some of the clusters from the training set are represented well (Figure 2c). While there are many out-of-distribution images, these still cluster to small regions of the entire distribution across all datasets.

Next we visualized the segmentation scores for the validation images, either the Cellpose scores (Figure 2d) or the difference in scores between Cellpose and Mediar (Figure 2e). This visualization points to two specific clusters where Cellpose and Mediar did better relative to each other: one cluster in the Tissuenet cloud where Cellpose did better, and one cluster in the DIC image region where Mediar did better. Thus, this more comprehensive analysis confirms the anecdotal observations we made inspecting single images above.

## Discussion

Here we have shown that a Cellpose model trained with default parameters outperforms the best result from the Neurips challenge, with no additional tuning. Thus, the results of this challenge were interpreted incorrectly by Ma et al [1]: transformers do not provide gains in performance for cellular segmentation, let alone the massive improvements advertised there. The analyses we presented above provide multiple cautionary tales, outlined below.

First, interpreting the results of public challenges can be difficult. Many methods tend to focus on complicated training protocols with many custom steps which help to overfit to a specific dataset with a specific metric, but do not necessarily result in generalizable improvements. Despite this, we believe that public challenges are beneficial for growing and engaging the community.

Second, in order to claim generalizable improvements in cellular segmentation one must perform interpretable analyses of the models. Ablation studies provide one approach to do this, and some ablations were indeed reported in the Mediar paper [2]. However, many of the results of the ablations were within a 1-2% performance range on a small validation set, which may not be enough to justify adding custom steps to the basic Cellpose pipeline.

Finally, we find that innovations in deep learning do not always translate to innovations in biological applications. In this case, transformers probably do not perform better because they are not trained on extremely large datasets such as those available in the computer vision field with hundreds of millions of images [14, 15]. Trained on “small” datasets of 1 million images, the performance of vision transformers is actually comparable to CNNs [16]. Since datasets of many millions of images are nearly impossible to label with human annotation, transformers are often trained on unlabeled data, which was also available for the Neurips challenge, but did not help any of the top teams [1]. In biology, it may just be impossible to collect datasets of millions of diverse images. For reference, putting together all the major currently available datasets, our “cyto3” model from Cellpose3 is trained on *∼* 8,000 images [10].

Given the results we have shown here as well as those we reported elsewhere [10], we conclude that transformers do not outperform Cellpose, and Cellpose remains the state-of-the-art in cellular segmentation.

## Acknowledgments

This research was funded by the Howard Hughes Medical Institute at the Janelia Research Campus.

## Author contributions

C.S. and M.P. designed the study, performed data analysis, and wrote the manuscript.

## Data availability

No new data was generated in this study. The Neurips challenge dataset is available at https://neurips22-cellseg.grand-challenge.org/dataset/. The ‘cyto2’ dataset is publicly available at https://www.cellpose.org/dataset, and the other datasets were generated and shared by other labs [7, 8, 17–23].

## Code availability

Cellpose3 was used to perform all analyses in the paper. The code and GUI are available at https://www.github.com/mouseland/cellpose. Scripts for recreating the analyses in the figures are available at https://github.com/MouseLand/cellpose/tree/main/paper/neurips.

## Methods

The Cellpose code library is implemented in Python 3 [24], using pytorch, numpy, scipy, numba, opencv, imagecodecs, tifffile, fastremap, and tqdm [25–33]. The figures were made using matplotlib and jupyter-notebook [34, 35].

### Datasets

For training the models for the challenge, we used the training images from the Neurips challenge dataset [1], and the training and testing images from the Cellpose ‘cyto’ dataset [3], the Databowl nucleus dataset [20, 21], the Livecell dataset [8], and the Omnipose dataset [7] (the four datasets that Mediar used). We included images from the test set of the external datasets, because Mediar also did. For the ‘cyto’ images that contained two channels (cells and nuclei), when training in RGB, these image channels were randomly assigned to two of the three color channels.

The Neurips challenge dataset consisted of 1,000 training images and 101 validation (“tuning”) images from various imaging modalities [1]. The fluorescent images in the dataset had cellular and nuclear markers which were placed in either the red, blue or green color channels. For evaluation, we did not use the last validation image, which was 8,000 by 10,000 pixels in size, because the evaluation for this image in the challenge was inaccurate: they used patches of size 2,000 by 2,000 to evaluate the F1 score, not requiring uniqueness for cell ids across patches and ignoring cells on the boundary of each patch [1].

For the t-SNE styles, we used the training images from the Neurips challenge dataset and from 9 publicly available datasets [7, 8, 11, 17–23], described in the Cellpose3 paper [10].

### Cellpose training and evaluation

#### Model architecture

As described in [3], the Cellpose model is a deep convolutional neural network with a U-net based architecture, consisting of four downsampling blocks and four upsampling blocks, each block containing four convolutional layers with residual connections [36, 37]. This network predicts XY flows and cell probabilities, which are turned into cell masks through an iterative dynamics process. The flow representations from Cellpose3 are used, which are a small update from Cellpose1 meant to address floating point errors in the calculations [10].

To compare to Mediar [2], we changed the backbone of Cellpose to a transformer-based segformer encoder (MIT-B5) [38] and a multi-scale attention decoder [39] (“Cellpose transformer”). As in Mediar, we used the implementation of the encoder and decoder from the segmentation models.pytorch github, and we used the pretrained imagenet weights provided for the encoder [40].

#### Training

For all training, we used the original loss function from the Cellpose paper [3]: the mean squared error between the XY flows from the ground-truth segmentation and the predicted XY flows, scaled by a factor of five, added to the binary cross-entropy between the ground-truth cell probability and the the predicted cell probability. For all images, each image channel was normalized such that 0 was set to the first percentile of the image intensity and 1 was the 99th percentile of the image intensity. The mean diameter for cells used to train the models was set to 30 pixels for all models.

To replicate the training performed in Ma et al [1], we trained Cellpose (“Cellpose impaired”) on the 1,000 challenge images converted to grayscale. The model was trained for 500 epochs with stochastic gradient descent with weight decay of 1e-5, momentum 0.9, and batch size 32. The learning rate increased linearly from 0 to 0.2 over the first 10 epochs, then decreased by factors of 2 every 10 epochs over the last 100 epochs. For the Cellpose-default and Cellpose-transformer models, we used three-channel inputs and the four additional datasets that Mediar used, using the same training parameters as in the Cellpose3 paper [10]. We used the AdamW optimizer with a weight decay of 1e-5 and batch size of 8 [12]. For training the Cellpose model with the standard CNN backbone, the learning rate increased linearly from 0 to 0.005 over the first 10 epochs, then decreased by factors of 2 every 10 epochs over the last 100 epochs. For training the Cellpose model with a transformer backbone, we used the same learning rate schedule, but the maximum learning rate was 0.0005 instead. The models were trained for 2,000 epochs where each epoch consisted of 800 training images, randomly sampled from the total training set of 7,692 images (1,000 challenge images + 6,692 training images from other datasets).

Each image from the other datasets had one or two channels. Images with only one channel were converted to grayscale, and images with two channels were converted to RGB with random channel assignment, placing cellular and nuclear channels in random positions. One channel images from the challenge dataset were converted to grayscale (same image in all three channels). The original Cellpose augmentations were used with training patches of size 224×224: random rotation, random flipping, and random resize with a scale factor between 0.75 and 1.25, such that 1.0 is equivalent to cell or nuclei diameters of 30 pixels [3]. We sampled the images from the Ma et al [1] training set with a probability of 75%. For the other four datasets (Cellpose, Livecell, Omnipose and DataScienceBowl-2018), we sampled images in equal proportion from each dataset (thus 6.25% probability from each dataset).

We trained a size model for the Cellpose-default model. For the size network training, we used the same defaults as in the Cellpose paper: 10 epochs of training images with random patches of size 512×512, and an L2 regularization constant of 1.0 [3]. Each epoch consisted of 800 images randomly sampled from the training set with the sampling probabilities described above.

#### Evaluation

We evaluated the main models by submitting to the “Testing” leaderboard of the Neurips challenge (Cellpose-default and Cellpose-transformer). We note that the current test images are different from the original challenge images: the authors state that they removed the easiest 50 images and added 30 new hard new images. We also evaluated on the Ma et al [1] validation set (called the “tuning set” in the paper and on the website). The tuning images were converted to grayscale for evaluation of the impaired Cellpose model and the Cellpose model from Ma et al [1] by setting the channels input in Cellpose to “[0,0]”. The flow error threshold (quality control step) was set to the default 0.4 and the cell probability threshold was set to the default 0. We turned on test-time augmentations during evaluation (as in Cellpose1 [3]) and used a tile overlap of 0.6, to match Mediar’s tile overlap size.

We ran the size model to estimate the cell diameters for the Cellpose-default and Cellpose-transformer models. Using the predictions of the size model, we resize each image to an average cell diameter of approximately 30 pixels, and we run Cellpose on this resized image. This is the approach introduced in the first Cellpose paper [3]. The running time of the size model is included in the reported run time per image for each of these models.

### Mediar evaluation

We used the “predict.py” script provided on the Mediar github to run the Mediar code on the validation images (using git commit 4339bca5a97c17282b2ffe4ac267e2999b7eab7f) [2]. We used the ensemble tta.json configuration file in the script, just changing the path to the dataset in the json to our local path. This ensures that Mediar uses test time augmentations and model ensembling.

### Quantification of segmentation quality

For quantification, we used the median F1 scores provided by the challenge website for test data, which exclude the boundary cells [1], or we used our own calculation of the F1 scores on validation data. Results were similar between using the evaluation code from and our own code, and also with or without the removal of boundary cells, so we opted not to remove such cells in our own calculation which is more standard practice.

To compute F1 scores, we proceed as in Cellpose 1, 2 and 3 by matching each predicted mask to the ground-truth mask that is most similar, as defined by the intersection over union metric (IoU) between the predicted and ground-truth. The F1 score for each image is defined using the true positives (matches with an IoU above a given threshold), false positives (predicted masks without matches), and false negatives (missed ground-truth masks):

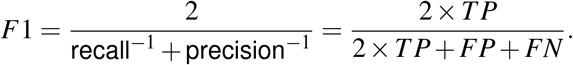

We compute the median F1 scores by taking the median across the 100 images in the validation set.

### Style vector embedding

We trained an additional Cellpose model on the training sets from 9 publicly available datasets and the training set from the challenge to obtain style vectors for two-dimensional t-SNE embedding like in the original Cellpose paper [3]. For image sampling during training, we set the challenge training dataset to 10% probability, and used all the other datasets from the Cellpose3 paper [10] in proportion to the sampling probabilites used there. We ran this Cellpose model on all the training images (9402 in total) and the validation images (100), extracting the style vectors (256-dimensional) from the output of the last downsampling layer of the network as previously described [3]. We then ran 2D t-SNE on the style vectors using the defaults in openTSNE: initializing with the top principal components scaled by 0.0001 and using a perplexity of 30 [13, 41]. The t-SNE embedding is visualized in Figure 2b.

